# SPIN: Inkjet-Driven Nanowell Workflow for Scalable and Sensitive Single-Cell Proteomics

**DOI:** 10.1101/2025.10.27.684859

**Authors:** Eric Cheng, Shuxin Chi, Huan Zhong, Robin Coope, Leonard J. Foster, Karen C. Cheung

## Abstract

Single-cell proteomics is emerging as a powerful approach to resolve cellular heterogeneity, yet sample processing remains challenging due to limited input material and the absence of protein amplification. We present a protocol centered on an image-guided, machine-learning-driven inkjet single-cell printer integrated with a dew-point-controlled nanowell chip to reduce loss, increase throughput, and improve reproducibility. The system dispenses single cells at >1 Hz into sealed nanoliter wells with minimal surface contact, virtually eliminating evaporation; a high-thermal-conductivity aluminum substrate and precise environmental control further ensure exceptional reproducibility. Relative to a commercial dispenser, the workflow yields significantly higher protein and peptide recovery without bias toward high-abundance species, delivering uniformly deep coverage. Biological pathway analysis emphasizes the robustness of this workflow, as there is a near 100% protein completeness detected among the enzymes in the Kreb cycle in both A549 and astrocytes, suggesting the consistency across all samples evaluated. This platform addresses core processing bottlenecks and enables reliable, scalable single-cell proteomics.

## Introduction

The Central Dogma suggests that there is a direct and obvious relationship between expression of an mRNA in a cell and that of its cognizant protein. However, measurements of mRNA and protein levels in the same samples, either bulk tissues or single cells, tend to reveal a poor correlation between the two, even when accounting for possible time lags (1-3). This incongruence suggests that regulatory mechanisms, such as post-transcriptional modifications, significantly obscure RNA-protein relationships (4). Elucidating the RNA-protein relationship more clearly requires quantifying both at the single cell level, and the challenge has been to profile proteins from single cells with as great a sensitivity and dynamic range as possible.

Proteins have historically been characterized by binding fluorescent or radio-labeled antibodies and measuring via imaging or blotting. Mass spectroscopy (MS) emerged this century as a non-targeted method to characterize a large fraction of all the proteins in tissue, leading to the concept of proteomics in parallel with the emergence of genomics and transcriptomics (5,6). Historical mass spectrometry (MS) methods have hitherto required millions of cells however, so are not able to capture uniqueness and intricacy in cellular subpopulations (7). This has been a driver for the development of single cell proteomics to better understand cellular function and regulation. Nucleic acid analyses, such as single-cell RNA sequencing, benefit from the ability to amplify the sample as well as to attach oligonucleotide bar-codes, enabling DNA or RNA from a large number of single cells to be individually prepared and pooled for sequencing (8). Without an equivalent method, the proteome’s large dynamic range causes signal losses from low abundance cell sub-populations, as bulk analysis can only construct an averaged proteome (9).

Despite these challenges, improvements in particularly the detection efficiency of mass-spectrometers, has put single cell analysis within reach (10,11). A cell, however, only has about 250 picograms of protein, and these molecules are prone to non-specific losses, particularly through binding to lab-ware. A goal therefore is to minimize manipulation and available surface area during sample preparation.

Techniques have emerged to overcome these difficulties. Groups have particularly focused on lowering sample preparation volumes to the nanoliter scale (12), preventing sample evaporation (13-15), and simplifying preparation steps (7,16). Others introduce workflows incorporating reagents, such as dimethyl sulfoxide (DMSO), to prevent sample evaporation and boost detection sensitivity (13,17,18). Together with several orders of magnitude gains in mass-spectrometer sensitivity, these advances have enabled compelling demonstrations of single-cell proteome coverage (11). However, obstacles still remain, as non-specific adsorption on plastic lab-ware during liquid-handling leads to sample loss, reducing overall protein identification rates and proteome depth (19,20). In addition, viscous detergents such as DMSO can introduce ion-suppression effects or carryover contamination, requiring extra cleanup steps to safeguard instrument performance and data quality (21,22). As a result, current single-cell proteomics workflows have predominantly focused on increasing the number of identified proteins, with limited emphasis on achieving high data reproducibility and depth of coverage for enabling comprehensive and robust pathway analysis (10). The combined challenges of low input material, technical variability in sample recovery, and missed detection of critical pathway components across individual cells limit the ability to reconstruct complete signaling cascades or metabolic pathways. High total protein identifications alone are insufficient if critical pathway components are inconsistently captured. Desirable features in single-cell proteomics workflows should, rather, highlight reproducibility across samples, consistent quantification of pathway-relevant proteins, and sufficient coverage to enable accurate biological interpretation. As a result, while individual protein quantification is increasingly feasible, system-level interpretations remain tentative. Here, we introduce Single-cell Proteomics in Isolated Nanowells (SPIN), through the use of the Isolatrix, an image-based inkjet single cell printer (23), and a nanowell substrate such as the Takara Bio Inc. Smartchip to enable a robust, miniaturized, one pot chemistry for single cell proteomics sample preparation. Individual nanowell contents are extracted by high-speed centrifugation through a bespoke funnel array that transfers each cell lysate into a separate MS-compatible container for analysis (24). The one-pot chemistry and centrifugation apparatus minimize liquid transfer steps, mitigating peptide loss from non-specific binding. The nanowell chip offers advantages in small (100nl) well volumes with high thermal conductivity for efficient lysis and digestion along with the capability to be non-intrusively sealed to facilitate efficient sample handling to maintain proteome coverage and depth. The Isolatrix uses a machine learning based cell dispensing algorithm which offers high throughput and high confidence in the single cell purity of the partitioned sample (23). The integrated substrate dew point control chiller mitigates sample loss to evaporation. Using A549 human lung adenocarcinoma cell line cells, we carried out comparison analyses to demonstrate that our workflow yields higher proteome coverage and can better resolve critical protein signaling pathways in comparison with a commercial cell dispenser dispensing into 384 well plates (25,26). By applying the SPIN workflow to primary astrocytes, it revealed distinct proteome signatures that underscore its ability to capture biologically meaningful heterogeneity.

## Results

### An integrated nanowell–inkjet platform for one-pot single-cell proteomics

The SPIN single-cell proteomics workflow is highlighted in Figure 1. The nanowell chip is an aluminum plate with a grid of 72×72 drilled holes, all coated with a polymer. Each well is 100 nL in volume and 460 μm in diameter with a well-to-well spacing of 542 μm. Cell samples were prepared (Supplementary Fig. 1) and loaded into the nanowells via the Isolatrix inkjet instrument which dispenses discrete droplets 160 pL in volume and is capable of addressing individual wells on the chip.

**Figure 1.**
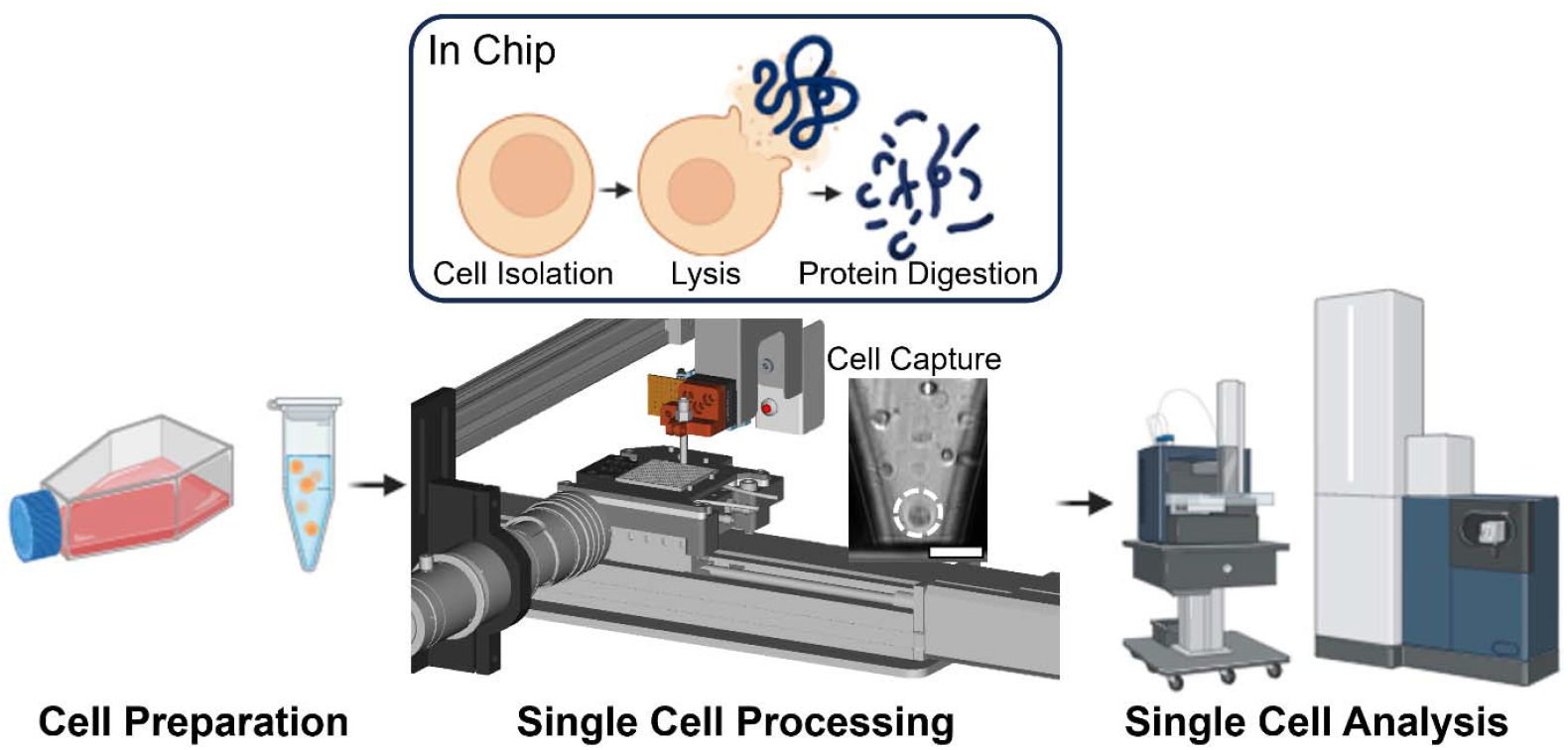
Schematic of the single-cell proteomics workflow using Isolatrix and nanowell chips for sample processing. Single cells are harvested and resuspended in 1x PBS. Using the Isolatrix system, cells are dispensed in nanowells for lysis and protein digestion. Samples are loaded onto Bruker NanoElute2 and timsTOF SCP. Scale bar in nozzle image represents 50 μm.

### Single-cell dispensing

Sample processing was performed in the nanowell chip as a one-pot chemistry (Figure 2a). Keeping the volume very low in this way maximizes enzymatic efficiency (27). The one-step, one-pot chemistry of the SPIN protocol facilitates efficient protein capture without introducing excessive contaminants on the mass spectrometer. The chip can also be sealed via a non-adhesive film (Bio-Rad Microseal A) to prevent sample cross-contamination and facilitate transport, centrifugation, mechanical lysis and incubation without the risk of introducing external proteins. To accommodate the film’s deformation into the well to create the seal, each well is filled to a maximum of 85 nL to prevent sample loss due to contact with the film.

**Figure 2.**
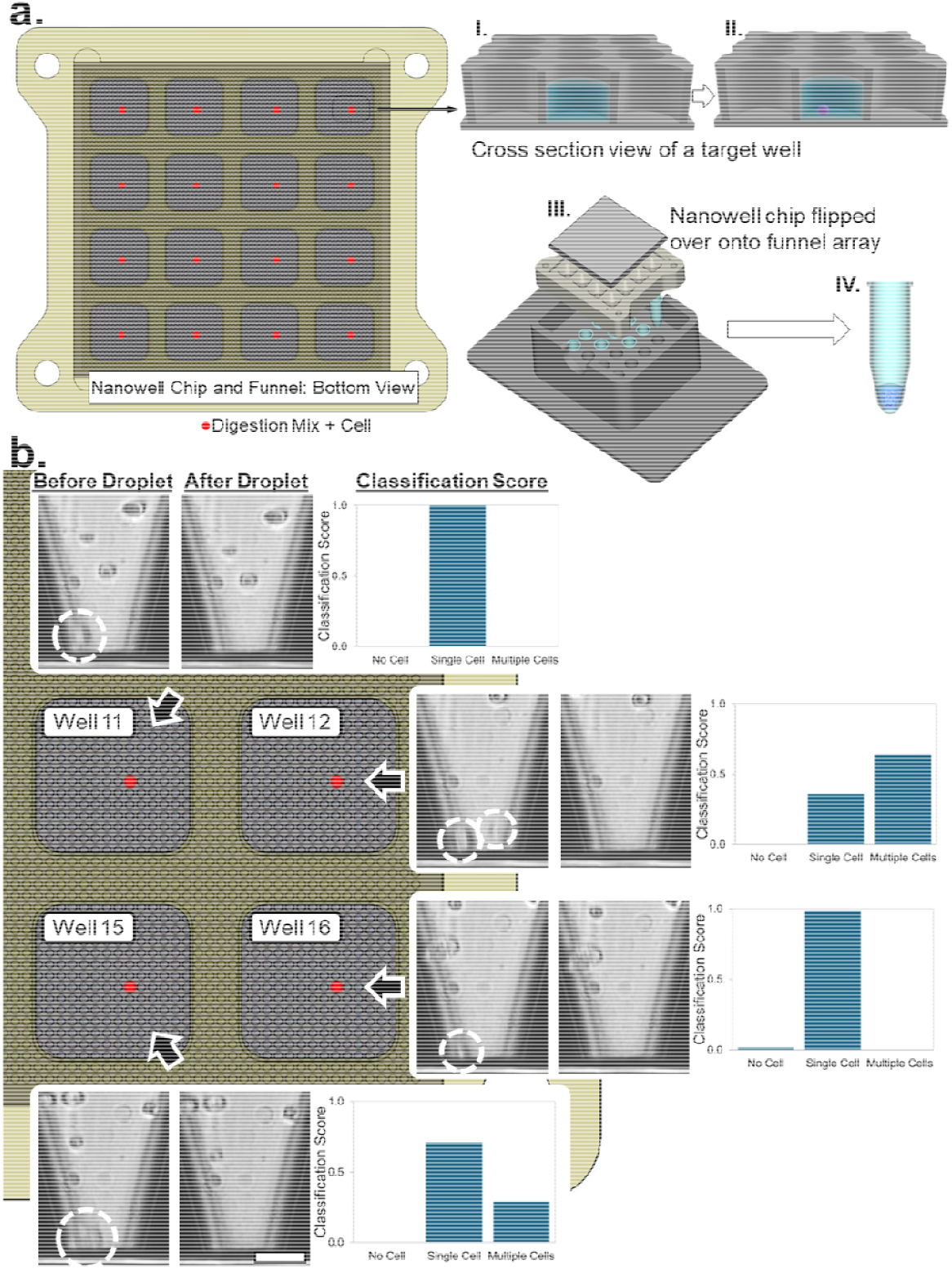
Diagram of the SPIN workflow. **(A)**.The nanowell chip (grey) is segmented into 16 regions, aligned over a funnel array (yellow). I. A single target well within each region pre-filled with a trypsin and DDM digestion mix. II. Single cell dispensing is performed on each pre-filled well. Lysis and digestion is performed on chip. III. The chip is placed in the funnel array where the cell dispensed well aligns with a designated funnel opening. To extract the contents of the nanowell, a spinning apparatus transfers the sample in solution to a collection tube via centrifugation at 3220 rcf. IV. The tubes are collected and placed in a custom holder for sampling on the mass spectrometer. **(B)**. Zoomed in view of the bottom right quadrant of a chip and funnel array. The red dot highlights the wells where the trypsin digestion mix and cells are dispensed. The nozzle image pairs of the cell dispensing droplet along with the instrument’s classification score is plotted. The dispensed cell, primary astrocytes from donor2, is highlighted. Wells containing multiple cells were omitted from analysis. Scale bar on nozzle image represents 50μm.

The funnel array segments the nanowell chip into 16 patches of 12×12 wells (Figure 2b). Each patch aligns with one of the funnel openings and one well is utilized within each patch for each experiment, so a chip could be used up to 144 times, isolating 16 cells each time. Trypsin was dispensed into these 16 wells using the Isolatrix inkjet instrument under dew point control via a Peltier cooling element positioned underneath the nanowell chip. Chilling of the substrate mitigated evaporation of the trypsin via an integrated dew point tracking unit (Supplementary Fig. 2). After reagent dispensing, the chip was sealed and stored in a −20°C freezer until cell dispensing.

Cells were then dispensed into the 16 trypsin-filled wells under substrate dew-point control. During dispensing, the instrument prints droplets over the target well until a cell-encapsulation event is detected or a timeout threshold is reached (a maximum of 20 droplets in this study). For each print run, four chips (16 wells per chip; 64 wells total) were prepared and processed in series. The instrument’s neural network analyzes the droplet image series and assigns a classification score to each of three classes: class 0 – empty droplet, class 1 – droplet containing a single cell, and class 2 – droplet containing multiple cells (Supplementary Fig. 3). The class with the highest classification score is taken as the final prediction of the cell-encapsulation event for each well.

**Figure 3.**
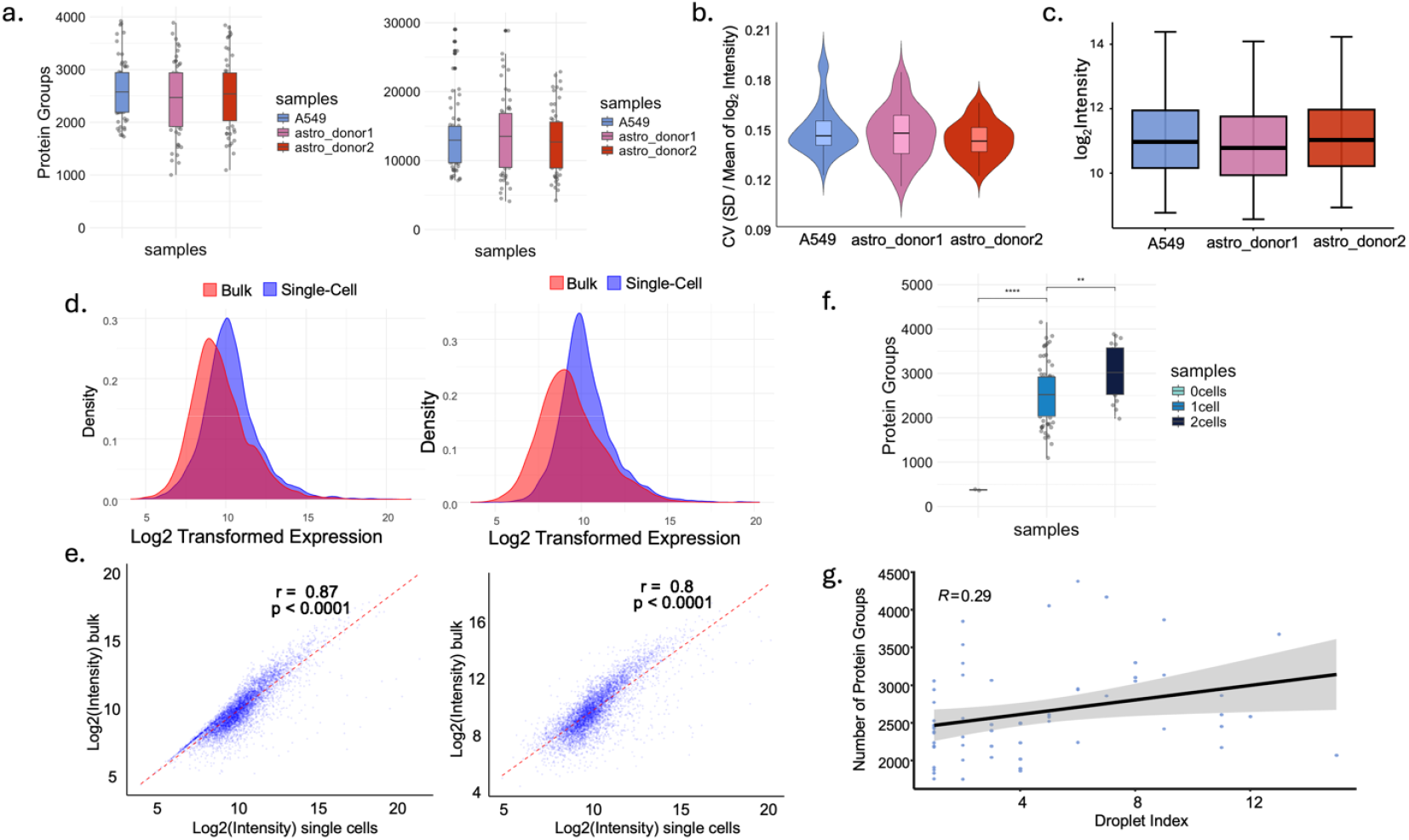
Single cell data obtained from the Isolatrix with the nanowell chip system. **(A)**. number of protein and peptide IDs identified for A549, astrocyte donor1 and astrocyte donor2; **(B)**. Coefficient of variation (CV) plot for all three cell types; **(C)**. log2 intensity of all proteins identified in three samples; **(D)**. Density plots showing the log2 transformed protein intensity for A549 (left) and astrocytes (right) compared to corresponding bulk samples; **(E)**. Correlations between bulk and single cell samples for A549 (left) and astrocytes (right); **(F)**. Number of protein groups identified with blank (0cells), 1 cell (1cell) and 2 cells (2cells); **(G)**. The correlation between the number of dispensed droplets in each well and the identified protein groups for the A549 cells with correlation coefficient (R=0.29) calculated.

As an example, when printing astrocytes, a chip with 16 target wells was completed in 9.3 s and yielded 14 wells containing a single cell and 2 wells containing multiple cells. One of the multiple-cell wells was misclassified as single-cell, resulting in an overall classification accuracy of 98.6% across all droplets dispensed into that chip. Across the full run of astrocyte donor1 (four chips, 64 wells total), 55 wells were correctly identified as containing isolated single cells, corresponding to an overall accuracy of 93.0%, with a total dispense time of 43.0 s. These performance metrics were consistent across all other cell types tested. All nozzle image series were manually inspected to validate the number of encapsulated cells in each well, and only wells confirmed to contain a single cell were included in the single-cell data analysis.

### Proteome coverage and data quality assessment

Three cell types, A549 (n = 66), primary astrocytes from a female donor (labeled as donor1, n = 58) and primary astrocytes from a male donor (labeled as donor2, n = 57), were collected with an average number of protein groups of 2672, 2560, and 2572 and peptide IDs of 12960, 13515, and 12692, respectively (Figure 3a). Data quality was assessed with protein dynamic range, coefficient of variation (CV) and correlation to bulk. A more-than-five-fold detection of protein log2 intensity was found for each cell type (Figure 3c), indicating good dynamic range. Low CV values around 0.15 were detected across all three cell types (Figure 3b). Comparative analyses were made between single cell and bulk data to evaluate their concordance and overall quality of the single cell data. Pearson Correlation Coefficient measured the linear correlations between single and bulk samples (Figure 3e). Both A549 (r = 0.871) and astrocyte (r = 0.799) had good correlation. Additionally, a comparison of density plots was made between bulk and single cell samples, with A549 samples showing a better overlap than that of astrocytes (Figure 3d). There was a significant increase in the total protein groups detected across 0 cells, 1 cell, and 2 cells from astrocyte donor2, further verifying the detection accuracy (Figure 3f). To assess if the variable droplet cell dispensing algorithm influenced the downstream protein digestion process, a negative binomial regression was run in SPSS on the A549 dataset (Figure 3g) which revealed that the variable dispense volume across the timeout conditions of 1-20 droplets was not a significant factor in the resulting protein group analysis (β = 0.012, SE = 0.037, p = 0.752). The small β indicates that the droplets had little effect on the resulting protein IDs and the p-value of 0.752 is not statistically significant, showing no correlation between dispensed volume and protein IDs in the tested range.

### Recovery in nanowells versus microplates

To further benchmark the relative robustness of the Isolatrix SPIN system, a comparison was made with the Tecan Uno Dispenser. Single cells of 200 cells/μl concentration were prepared as required by the Tecan Uno Dispenser and dispensed into 384 well plates with a total volume of 1.5 µl for sample preparation. As shown in Figure 4a, the nanowell chip provided an increase of protein detection in all samples, especially primary astrocytes with a 23.5% increase in donor1 and 18.5% increase for donor2. Quantitative coverage was assessed using protein completeness across all single cells from each system (Figure 4b). Comparatively, Isolatrix data had 500 more proteins in the 80-100% completeness range across all samples, demonstrating better protein recovery and higher data quality. In the A549 sample, 79.2% of the identified proteins using the SPIN protocol had better completeness compared to the samples prepared in well plates. Additionally, 765 proteins were identified within the SPIN samples which were completely missing in the cells prepared in well plates. Protein correlation was examined between Isolatrix and Tecan Uno (Figure 4c). Overall, data from these two instruments had a 70-80% correlation, with slightly better correlation in A549 than primary astrocytes. The same A549 samples were injected on both the timsTOF SCP and timsTOF Ultra2 (Supplementary Fig.4). As expected, timsTOF Ultra2 detected 25% more protein groups. CV was calculated for both samples from timsTOF SCP and Ultra2, showing low variabilities between single cells, regardless of the mass spectrometer used.

**Figure 4.**
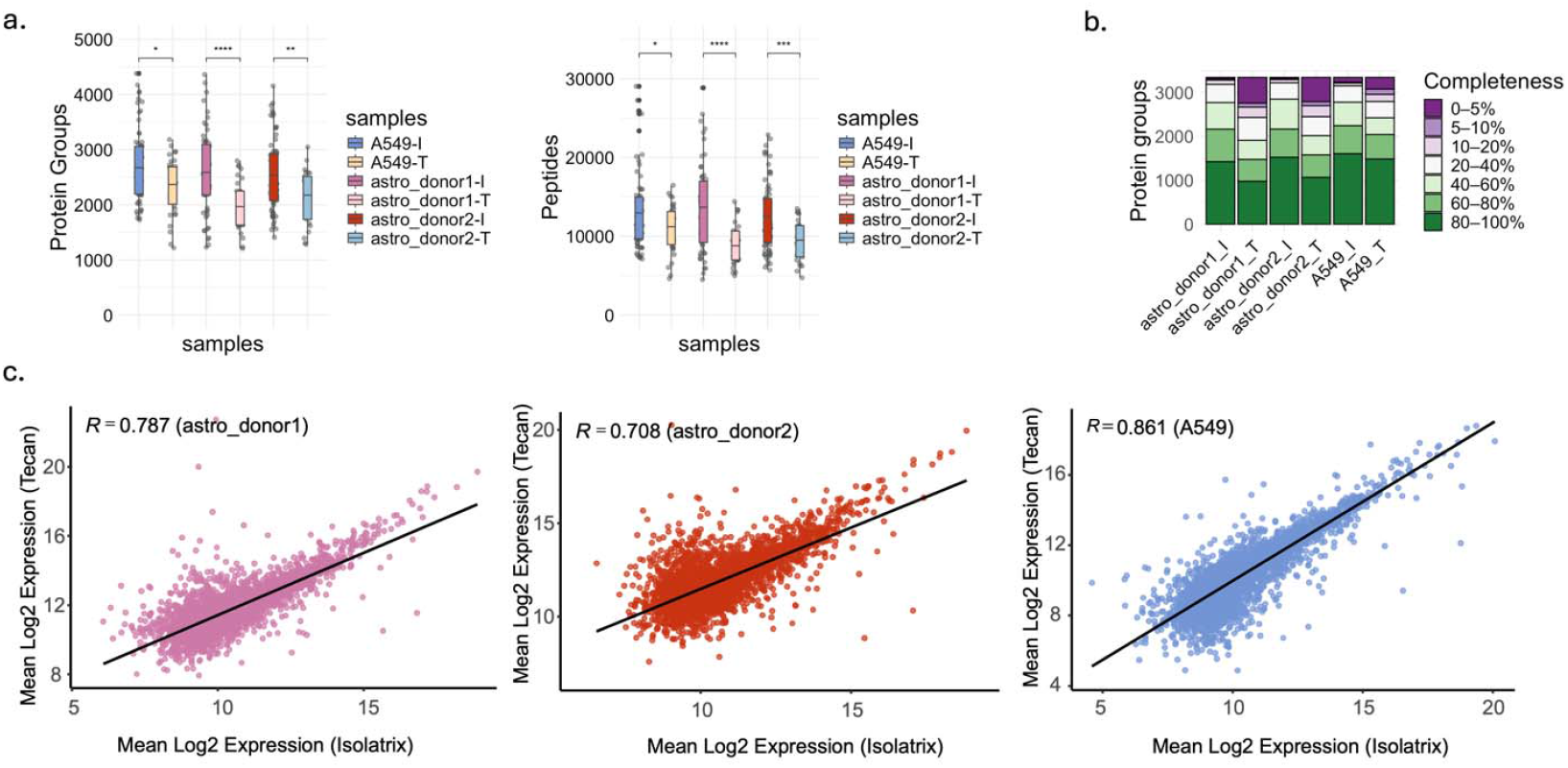
Comparison between Isolatrix and Tecan Uno Dispenser. **(A)**. Protein groups and peptides identified using Isolatrix (I) and nanowell chip system compared to Tecan Uno (T) with 384 well plates. **(B)**. Quantitative comparison between Isolatrix and Tecan Uno results for all three cell types. Each bin represents the completeness of protein coverage in percentage. **(C)**. Correlations between log2 protein intensity for A549, astrocyte donor1, and astrocyte donor2 from Isolatrix vs. Tecan Uno.

The benefit of using the SPIN method was further examined by comparing data from nanowell chip and well plates using the Isolatrix. The protein and peptide detection showed a 17.6% and 26.4% increase, respectively, in identification in the A549 samples prepared in the Smartchip compared to the same samples prepared within the 384-well plate (Figure 5a). Both a Venn diagram and a Principal Component Analysis (PCA) were made to examine the similarity and difference between chip and well plate data. There were 862 unique proteins identified from the chip workflow and 228 proteins from the plate wells (Figure 5b). The PCA plot showed a partial overlap, however, the PC1 and PC2 percentages are low, showing a small variation between chip and plate well (Figure 5c). The data quality was assessed through a protein sub-localization plot (Figure 5d). The percent of proteins identified in eight different cellular compartments were evaluated. Samples analyzed from nanowell chips displayed a higher protein coverage in every cellular compartment compared to that from well plates.

**Figure 5.**
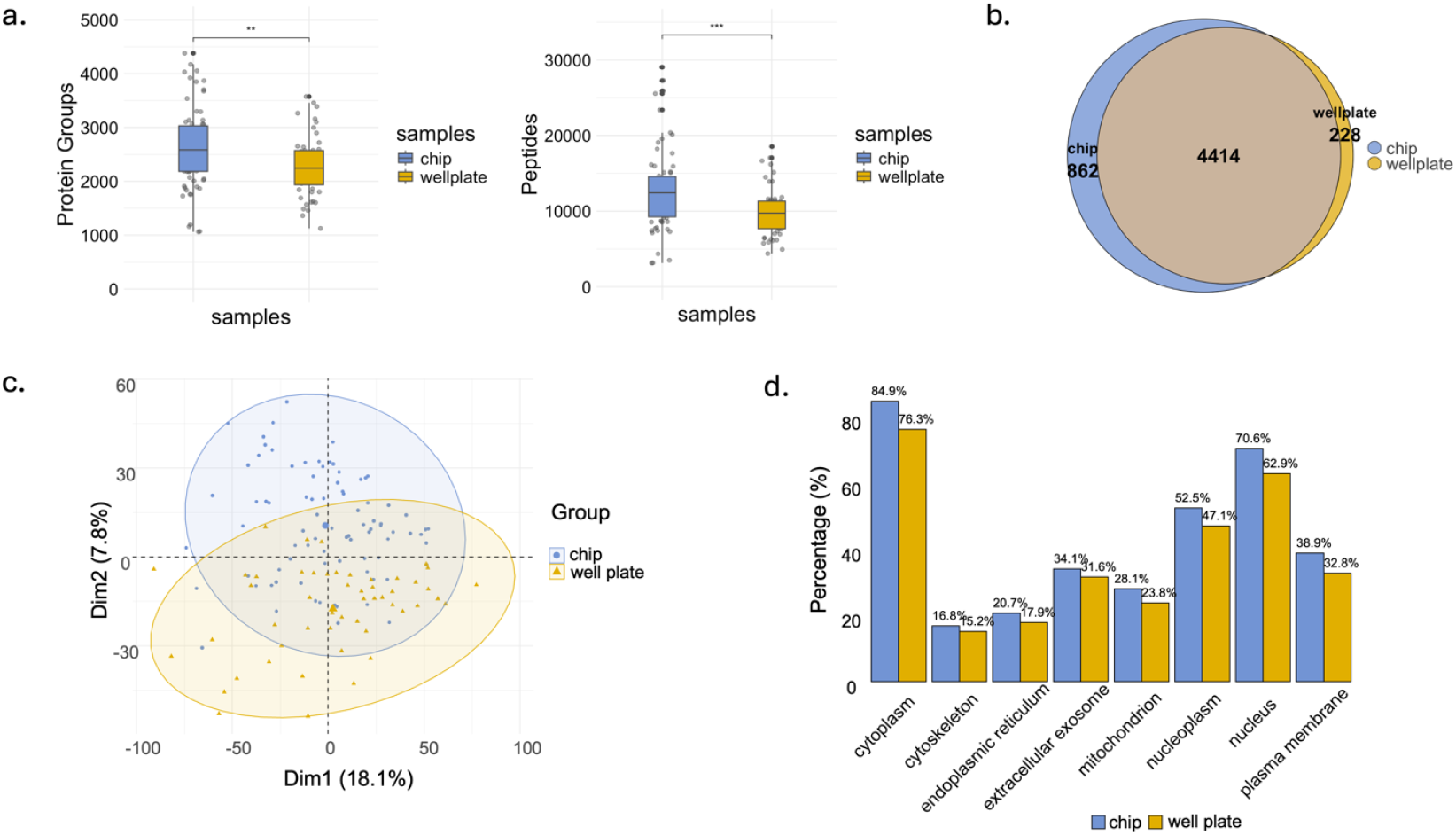
Comparison between samples prepared with nanowell chips and 384-well plates using the Isolatrix. **(A)**. protein and peptide IDs identified using nanowell chip (n = 66) and 384-well plates (n = 51). **(B)**. Venn diagram presenting the overlap and differences in the proteins identified between nanowell chips and well plate wells. c. Principal Component Analysis (PCA) between nanowell chips and well plates. d. Protein sub-localization comparison between nanowell chips and well plates.

### Dissecting cellular heterogeneity and pathway completeness

The biological context of the single cell proteome data was also investigated. Uniform manifold approximation and projection (UMAP) of all three cell types were presented with A549 having more distinct clustering than the primary astrocytes (Figure 6a). An astrocyte marker, GFAP (28), was highlighted in UMAP, which was exclusively presented in primary astrocytes (Figure 6b). Protein markers for each cell type were obtained through differential analysis with limma on the bulk dataset (Supplementary Fig. 5). The presence and intensity of these protein markers were measured in single cell data (Figure 6c, Supplementary Fig. 6). In particular, astrocyte protein markers, such as GFAP, FLNC, and PRDX2, had good sample completeness in astrocytes with high intensity; A549 markers, such as MYL, KRT, AGR2, and MISP, had good coverage and intensity in A549; and the rest had higher intensity in the corresponding cell types, but less perfect sample completeness.

**Figure 6.**
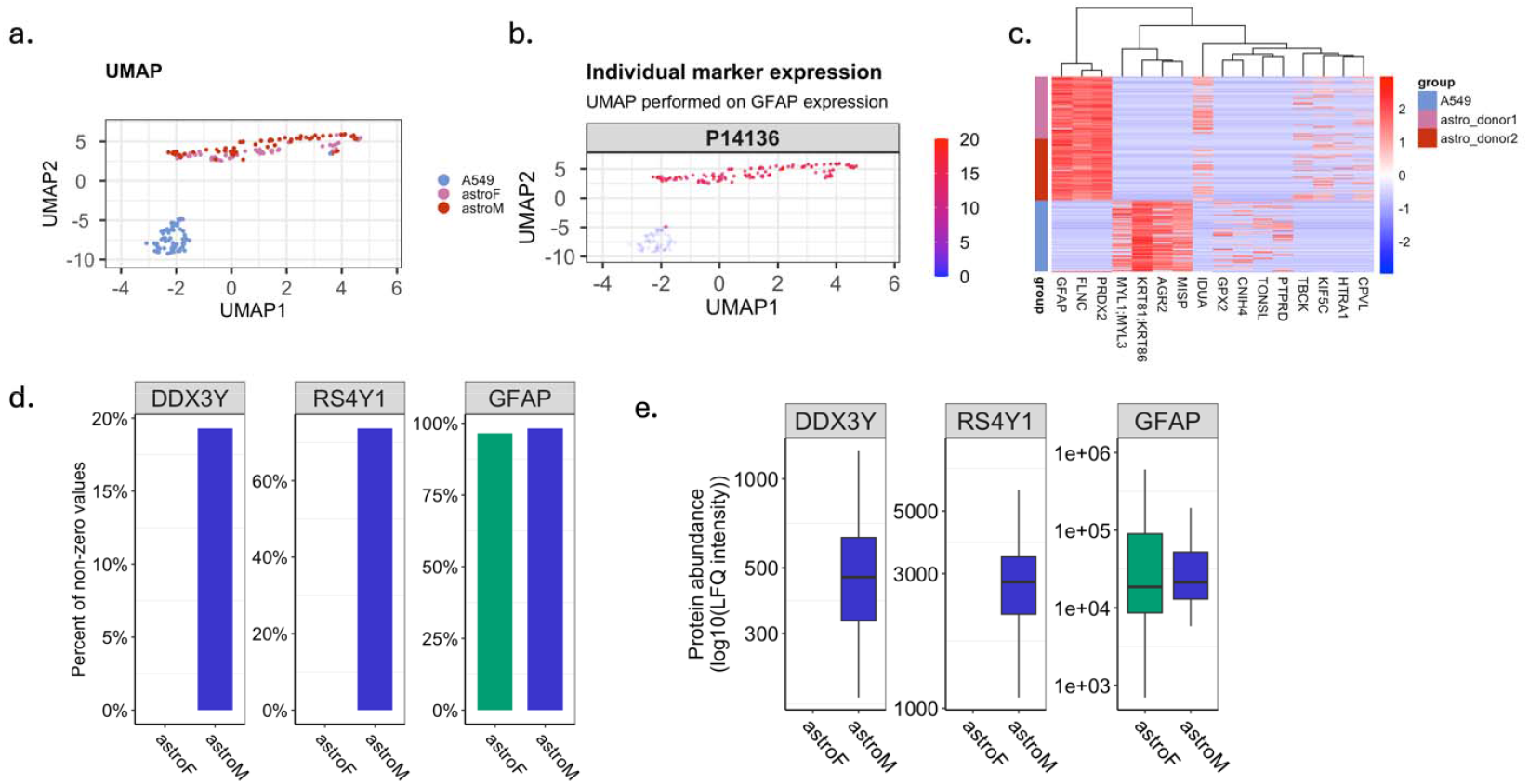
Bioinformatic analysis of A549 and primary astrocytes. **(A)**. PCA of A549, astrocyte donor1, and astrocyte donor2. **(B)**. UMAP showing the overall clustering of A549, astrocyte donor1, and astrocyte donor2 (top) and GFAP expression of all cells (bottom). **(C)**. Heatmap displaying the marker proteins for both A549 and astrocytes. **(D)**. Sex specific protein markers for male astrocytes (donor2), compared to female (donor1).

The primary astrocytes were collected from donors of the opposite sex (Figure 6d). The sex variability was also evaluated. DDX3Y and RS4Y1 are both Y-chromosome proteins, which were present in the male donor (donor2) only, validating workflow precision. GFAP was used as a reference.

Pathway analysis was conducted to explore potential clinical applications with biological meaning (Figure 7). The protein completeness of data collected from the nanowell chip and well plate was plotted. The Krebs cycle, a core pathway in cells, was examined across all three cell types. As shown by the Radar plot (Figure 7a), most proteins involved had near 100% completeness, with one exception - succinate dehydrogenase in primary astrocytes (Supplementary Fig. 7). We also examined the mTOR pathway, where we observed that the chip (orange) had generally higher data completeness compared to well plate (pink), which further validated the results earlier (Figure 7b).

**Figure 7.**
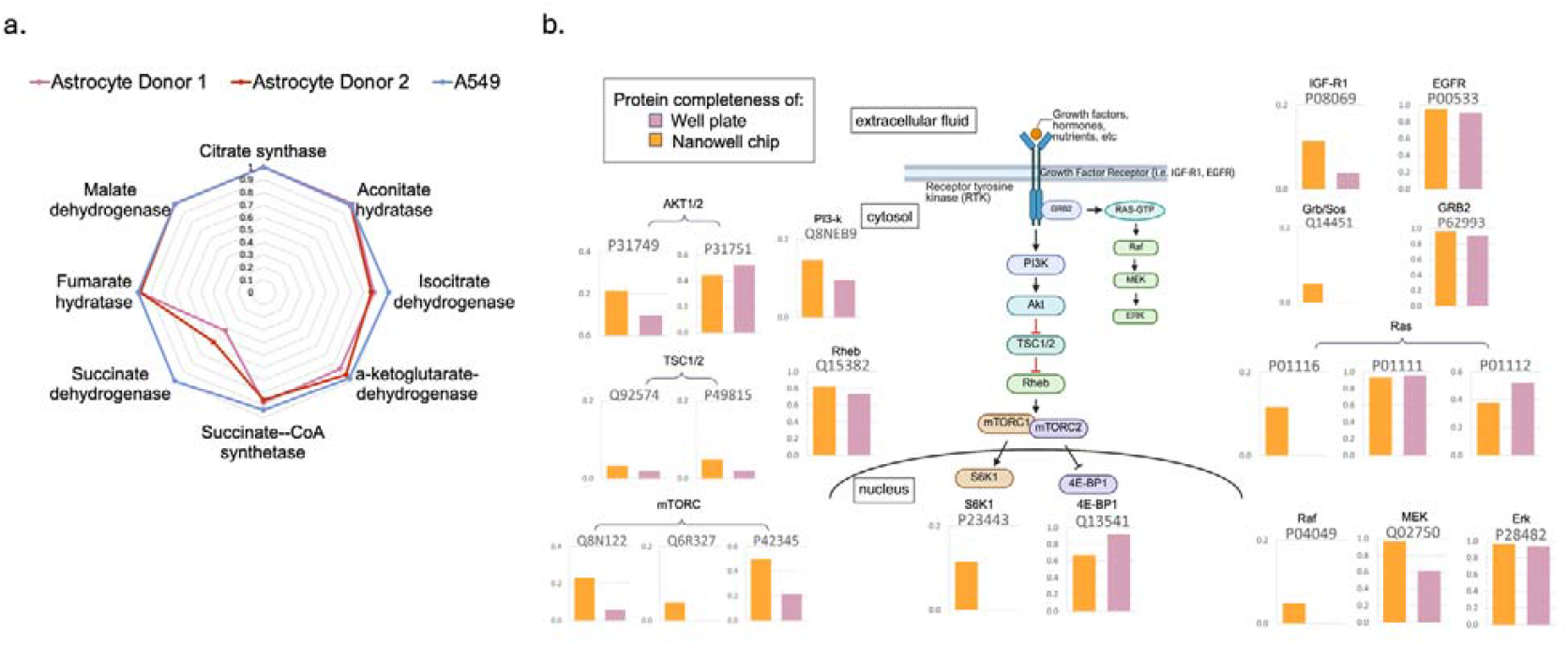
Pathway analysis of A549 and primary astrocytes. **(A)**. Data completeness of proteins involved in the Krebs cycle pathway. **(B)**. Comparison between the nanowell chip and well plate well data quantitative coverage for the mTOR pathway in A549.

## Discussion

A goal of single cell proteomics is to enable biological insights, which means data reproducibility is important. We therefore set out to build a method to get good protein counts and high reproducibility, starting from the premise that low lysis and trypsin digest volume would be of benefit in peptide extraction and digestion through improved reaction efficiencies. Here, we present an integrated system using the Isolatrix, an image-based inkjet single-cell dispensing instrument coupled with the nanowell chip for our SPIN sample preparation protocol. Our results bear our hypothesis out, as we saw consistent protein quantification and better protein completeness in both A549 and primary astrocyte data.

Quality assessment matrices, such as CV, dynamic range, and comparison to bulk samples, were used to elucidate the high reproducibility and unbiased proteome coverage using the Isolatrix system. Importantly, the improved sensitivity extended beyond high protein detection, reflecting more complete and unbiased proteome coverage. The advantages of the SPIN system, using the Isolatrix with nanowells were highlighted through direct comparisons against the Tecan Uno using its smallest supported (384) well-plate format and the Isolatrix spotting into 384 well plates. The protein quantitative coverage assessment demonstrated high data quality beyond just the number of proteins identified. As shown in Fig. 4b, the Isolatrix data displayed higher reproducibility and consistency across samples, proving higher reliability and data quality enabling better downstream biological investigations. The aluminum construction of the nanowell chip allows for controlled regulation of the substrate’s temperature in order to mitigate evaporation during dispensing and facilitate incubation during protein digestion. The small reaction volumes allow for efficient enzymatic reaction kinetics without the use of complex chemistries or the need for sample cleanup, enabling sufficient proteome coverage, sensitivity, and reproducibility required for downstream biological analysis.

Biological validation using UMAP, and GFAP astrocyte marker expression plots showed a clear clustering difference between A549 and primary astrocytes, highlighting the SPIN workflow’s ability to resolve meaningful cellular diversity. The mTOR pathway, as a central regulatory axis for cellular metabolism, growth, and homeostasis, is essential in diverse tissues (e.g., A549 lung epithelial cells) and is frequently dysregulated in diseases ranging from metabolic disorders to cancer (29). Its role in oncogenesis and therapeutic resistance has made it a major focus of targeted cancer therapies, leading to extensive research into mTOR inhibitors (30). Leveraging the nanowell’s finely controlled experimental conditions, our SPIN workflow achieved significantly higher completeness for mTOR pathway proteins compared to conventional well-plate preparations, ensuring robust detection of both upstream regulators (e.g., PI3-K) and downstream effectors (e.g., S6K, mTORC). Similarly, near-complete coverage of key Krebs cycle enzymes underscores this platform’s capacity to profile central metabolic networks at single-cell resolution. The only noticeable exception is succinate dehydrogenase, likely due to its membrane-bound nature rather than a limitation of the system (31). Together, these analyses highlight the SPIN workflow’s capability to deliver comprehensive, unbiased pathway profiling in individual cells.

Very small reagent volumes enabled by the Isolatrix and nanowell chip offers key advantages for proteomics. Compared to glass slides and microfluidic devices (32,33), the wells physically separate the samples to reduce potential cross-contamination. The non-adhesive, silicone pressure seal allows truly contamination-free sample handling, while an integrated real-time dew-point-controlled substrate chiller prevents evaporation and ensures volumetric precision, this avoids the introduction of exogenous reagents, such as mineral oil, that may interfere with mass spectrometry. By performing the one pot chemistry processes entirely on chip and only extracting peptides via the funnel array, we minimize sample loss, obviate the need for a cleanup step, further increasing capture efficiency.

The Isolatrix printer demonstrated compatibility with a number of cell types without introducing biases such as by cell size. The instrument’s neural network classifier and the print run data availability allows for manual inspection of dispensing events, providing high confidence that the captured content originates from a genuine single cell. The instrument achieves a throughput of greater than one cell per second and was only limited by the number of addressable wells per chip as dictated by the funnel array design.

The ability to use a small volume of comparatively concentrated trypsin in the nanowells enhances reaction kinetics, shortening trypsin digestion time and eliminating the need to overload enzymes, which can introduce contaminants into the mass spectrometer (34). Each nanowell supports reactions in miniaturized volumes while the thermally conductive wells enhance lysis and enzymatic digestion efficiency. Once processed, samples are extracted via high-speed centrifugation through a custom-machined funnel, ensuring minimal contamination and sample loss, improving overall protein recovery.

In this work, the array of 16 funnels limited the number of target cells per chip. To address this limitation, four chips were prepared in series to improve yield per run. The design and fabrication of a new denser funnel array to complement the existing setup can greatly increase the density of the addressable wells on chip by creating a setup which extracts directly into denser LC compatible 384-well plates. Additionally, bespoke nanowell arrays purpose-machined for proteomics work can further optimize the process, such as by enabling direct aspiration from the substrate into the LC without the need for transferring into a tube for sampling (35,36). While a number of inkjet-based cell dispensers may be appropriate for the SPIN workflow, the Isolatrix offers advantages in data traceability and scalability. Recent advancements in liquid chromatography columns have demonstrated significantly higher sample injection rates, establishing a foundation for high-throughput single-cell proteomics (37-39). With a higher utilizable density substrate coupled with the Isolatrix (23), the SPIN workflow enables practical scaling to thousands of single cells per experiment, addressing potential limitations in dispensing speed observed in current workflows (18). This allows rapid cell isolation within minutes, facilitating comprehensive and representative profiling of cellular proteomes, positioning the platform to meet the demands of next-generation, high-throughput mass spectrometry systems. Indeed, this workflow easily meets the scaling needs for current single-cell proteomics needs targeting a few hundred cells per condition. At this scale, LC-MS/MS time and data analysis steps are very much rate-limiting. As these latter steps become faster with time, the Isolatrix and SPIN workflow will continue to be future-proofed. Even more significant, in our opinion, is the increased proteome coverage enabled by the low and reproducible sample volumes that the Isolatrix delivers.

In summary, the SPIN workflow delivers a miniaturized, high-throughput, contamination-resistant, and cleanup-free solution for single-cell proteomics, enabling reproducible, in-depth proteome profiling. Looking ahead, this platform can be expanded to integrate with automated liquid-handling systems and high-density nanowell arrays, achieving truly high-throughput single-cell proteomics analysis. Coupling our miniaturized chemistry with emerging spatial proteomics and multi-omics integration strategies will further enhance our understanding of the molecular architecture of tissues, paving the way for unprecedented insights into cell-cell interactions and microenvironmental influences.

## Materials and Methods

### Cell culture

#### A549

A549 human lung adenocarcinoma cell line cells (ATCC, CCL-185) were cultured at 37°C with 5% CO_2_ with humidity control and maintained with Dulbecco’s Modified Eagle’s Medium (DMEM) containing 10% Fetal Bovine Serum (FBS), 1% penicillin/streptomycin, and 1% sodium pyruvate. The cells were subcultured or harvested at 80% confluency. To harvest cells, 1 mL 0.25% trypsin/EDTA solution was added for detachment. The collected pellet was washed three times with 1x Phosphate Buffered Saline (PBS).

#### Primary human astrocytes

Primary male (donor lot#: 31978) and female astrocytes (donor lot#: 36204) from the human cerebral cortex (ScienCell) were cultured at 37°C with 5% CO_2_ with humidity control. Astrocyte medium was also purchased from ScienCell including 2% FBS, 1% Astrocyte Growth Solution (AGS) and 1% penicillin/streptomycin. The cells were subcultured every 4 days and the passage numbers were controlled between passage 3 to 5 for experiments. Cell harvest was done by adding 10 mL 0.025% trypsin/EDTA and washing three times with 1x PBS.

### Dew Point Control in Nanowell Chip

A custom substrate chiller uses a Peltier element mounted directly beneath the nanowell chip (Takara Bio Smartchip) to actively regulate its temperature. An integrated temperature and humidity sensor (Telaire T9602) continuously monitors the ambient conditions, and the system uses these measurements to calculate a dynamic setpoint that cools the substrate just below the dew point, thereby minimizing sample evaporation.The Peltier maintains setpoint to ±0.1□via an integrated controller (Oven Industries 5R7-573) and a custom liquid cooling channel loop regulates the hot side of the element. Setpoint temperature is set to the calculated dew point minus 2□. The dew point temperature is updated every 10 seconds. All printing operations were performed at dew points above 6□.

### Single cell dispensing with Isolatrix

Cell dispensing and reagent printing into the nanowell chip was performed on an inkjet-based system developed in house and described in previous work (23). The target wells on the chip were predefined and reagents were dispensed into under dew point control. A total of 500 droplets of the digestion mix, each with a volume of 160 pL, were dispensed into each well, resulting in an overall deposition of 80 nL into the 100 nL well. Post reagent dispensing into each of the 16 wells, the Smartchip was sealed (BioRad MicroSeal A) and stored at −20□until cell dispensing.

For cell dispensing, the input cell concentration was measured with a cell counter (Logos Biosystems Luna II) and diluted to a target input cell concentration of 600 cells/μL. The timeout condition was set to a maximum of 20 droplets. Where the well was skipped after 20 consecutive empty droplets into the target.

After cell dispensing, the chip was sealed, and the cells were mechanically lysed through a freeze-thaw cycle in a −80°C freezer for 10 minutes and then incubated in a thermocycler (BioRad MJ-Mini) under a heated lid at 37°C for 90 minutes for protein digestion. Digested peptides were pulled into LC compatible 100 µL microcentrifuge tubes (GeneBio Systems Inc. PCRS001) using a funnel array at 3220 rcf for 5 min. The microcentrifuge tubes were pre-filled using a positive displacement contactless liquid handler (SPT Labtech Dragonfly) with 1.4 µL of a 0.015% (n-Dodecyl β-D-maltoside) DDM solution to increase the sample volume for LC uptake (40). The tubes were then placed in a bespoke 3D printed holder, mimicking the layout of a 96 well plate for LC sampling.

### Single cell dispensing with Tecan Uno

Cell concentration was measured with Countess II and diluted to a target input cell concentration of 200 cells/μl for dispensing. Single cell isolation was achieved using a Tecan Uno Single Cell Dispenser, where each cell was isolated into Eppendorf twin.tec PCR lowbind 384-well plates containing 0.5 µL LC/MS graded water. Based on the measured average cell size from Countess II, the Tecan Uno was set to dispense using cell size medium (15-17 μm) for A549 and large (18-20 μm) for primary astrocytes. The cells went through one cycle of freezing and thawing for lysis. Using the same dispenser, 1 µL of 1 ng LC/MS grade Trypsin/LysC in 50 mM triethylammonium bicarbonate (TEAB) with 0.03% DDM was introduced to each cell for protein digestion. The digestion occured at 37° for 1.5 hours with humidity control. The samples were stored at −70° until ready for MS.

### LC-MS/MS analysis

#### Liquid chromatography (LC)

Single cell samples were injected and separated on-line using NanoElute 2 UHPLC system (Bruker Daltonics) with Aurora Series Gen3 (CSI) analytical column, (25 cm x 75 μm 1.6 μm C18 120Å, with CSI fitting; Ion Opticks, Parkville, Victoria, Australia). The analytical column was heated to 50°C using a column toaster M (Bruker Daltonics). Buffer A consisted of 0.1% aqueous formic acid and 0.5% acetonitrile in water, and buffer B consisted of 0.1% aqueous formic acid and 0.5% water in acetonitrile. Before each run, the analytical column was conditioned with 4 column volumes of buffer A. The analysis was performed at 0.30 μL/min flow rate. The samples were run with a 30 minute gradient (from 2% B to 12% B over 15 minutes, then to 33% B from 15 to 30 minutes, then to 95% B over 0.5 minutes, held at 95% B for 7.72 minutes) for Isolatrix data and a 15 minute gradient for the Tecan data (from 2% B to 12% B over 7.5 minutes, then to 33% B from 7.5 to 15 minutes, then to 95% B over 0.5 minutes, held at 95% B for 5 minutes).

#### Mass spectrometer (MS) - timsTOF SCP

NanoElute 2 LC was coupled to a Trapped Ion Mobility - timsTOF SCP (Bruker Daltonics). The Captive Spray ionization source was operated at 1700 V capillary voltage, 3 L/min drying gas and 200°C drying temperature. During analysis, the timsTOF SCP was operated with Parallel Accumulation-Serial Fragmentation (PASEF) scan mode for DIA acquisition. The MS spectra were collected in positive mode from m/z 100 Th to m/z 1700 Th, and from ion mobility range (1/ K0) 0.7 V*s/cm2 to 1.3 V*s/cm2. The TIMS was operated with equal ramp and accumulation time of 100 ms (100% duty cycle). For each TIMS cycle, 11 dia-pasef scans were used, each with 3-4 steps. A total of 36 dia-pasef windows span from m/z 299.5 Th to m/z 1200.5 Th, and from ion mobility range (1/ K0) 0.7 V*s/cm2 to 1.3 V*s/cm2 with an overlap of m/z 1 Th between two neighboring windows. The collision energy was ramped linearly as a function of mobility value from 20eV at 1/k0 = 0.6 V·s/cm2 to 65eV at 1/k0 = 1.6 V·s/cm2.

#### Mass spectrometer (MS) - timsTOF Ultra2

timsTOF Ultra2 (Bruker Daltonics) system with the same LC set up was used. The Captive Spray ionization source was operated at 1600 V capillary voltage, 3 L/min drying gas and 200°C drying temperature. During analysis, the timsTOF Ultra2 was operated with Parallel Accumulation-Serial Fragmentation (PASEF) scan mode for DIA acquisition. The MS spectra were collected in positive mode from m/z 100 Th to m/z 1700 Th, and from ion mobility range (1/ K0) 0.66 V*s/cm2 to 1.42V*s/cm2. The TIMS was operated with equal ramp and accumulation time of 150 ms (100% duty cycle). For each TIMS cycle, 8 dia-pasef scans were used, each with 3 steps. A total of 24 dia-pasef windows span from m/z 400 Th to m/z 1000 Th, and from ion mobility range (1/ K0) 0.64 V*s/cm2 to 1.37 V*s/cm2 with an overlap of m/z 1 Th between two neighboring windows. The collision energy was ramped linearly as a function of mobility value from 20eV at 1/k0 = 0.6 V·s/cm2 to 59eV at 1/k0 = 1.60 V·s/cm2.

### Data analysis

All acquired data was searched on DIA-NN (version 1.8.1) against a database consisting of the FASTA and common contaminants, as well as a bulk library generated from 10 ng injection of the sample on timsTOF SCP (41). Fasta digest and deep learning-based spectra, RTs and IMs prediction were enabled using trypsin/P protease specificity and 1 missed cleavage. Other search parameters include N-terminal M excision. Peptide length ranged 7-30, precursor charge ranged 2-4, precursor m/z ranged 300-1200, and fragment ion m/z ranged 200-1800. Precursor FDR was set to 1%, with 0 for settings ‘mass accuracy’, ‘MS1 accuracy’ and ‘scan window’. Settings ‘heuristic protein inference’, ‘use isotopologues’, ‘match between run (MBR)’, and ‘no shared spectra’ were all enabled. ‘Gene’ was chosen for protein inference parameter along with ‘double-pass mode’ for neural network classifier. Robust LC (high precision) was used for quantification strategy, RT-dependent mode for cross-run normalization, and smart profiling mode for library generation.

### Bioinformatics

Quantified protein intensities were log_2_-transformed for statistical analysis and visualization. Bulk samples were assessed using limma (42) to obtain the specific protein markers associated to each cell type. To evaluate data quality and consistency of the single cell data, we performed multiple comparative analyses across sample types, donors, and platform technologies. Protein and peptide identification depth was compared across A549 cells, astrocyte donor1, and astrocyte donor2 using different sample processing platforms, including the Smartchip systems and Tecan Uno (384-well plate format). Protein coverage completeness per sample was binned by quantification frequency to compare data depth.

To assess reproducibility, Pearson correlation coefficients were calculated between average protein intensities derived from single-cell runs using Isolatrix versus Tecan Uno systems across all three cell types. Protein detection overlap was further visualized using Venn diagrams. Principal component analysis (PCA) was performed to examine global proteome differences between chip- and well-based data, and between cell types. Subcellular localization patterns were annotated using UniProt terms and summarized by organelle compartment to assess proteome coverage bias.

To explore biological identity and functional relevance, we visualized cell-type-specific markers using UMAP from scDataviz (43), heatmaps, and boxplots. These included known astrocyte markers such as GFAP, and sex-specific markers (e.g., DDX3Y for male donor). Completeness of coverage for metabolic proteins (e.g., TCA cycle) and signaling proteins (e.g., mTOR pathway) was quantified and compared between Smartchip and well plate formats.

All analyses were conducted in R (v4.4.1), using packages including tidyverse (44), ComplexHeatmap (45,46), FactoMineR (47), and ggplot2 (48). Pathway annotations were curated from KEGG (49-51) and UniProt (52) databases.

## Acknowledgments

We thank Erin Pleasance for useful discussions. We thank Peter Lansdorp for the funnel array. The funnel array setup was made at the BC Cancer Joint Engineering Centre by Scott Young.

## Author contributions

Conceptualization: EC, SC, LJF, KCC

Methodology: EC, SC

Investigation: EC, SC

Visualization: EC, SC, HZ

Supervision: LJF, KCC

Writing—original draft: EC, SC

Writing—review & editing: EC, SC, HZ, RC, LJF, KCC

## Competing interests

KCC and EC are inventors on a patent application covering elements of the dispensing technology (CA3149667/US20220155331A1/EP4010681A4/CN114341617A/WO2021022374A1).

## Data and materials availability

All data are available in the main text or the supplementary materials.

## Supplementary Materials

### Supplementary Text

The supplementary section includes extended technical descriptions of our SPIN set-up and additional data analysis used to support the discussion.

**Fig. S1.**
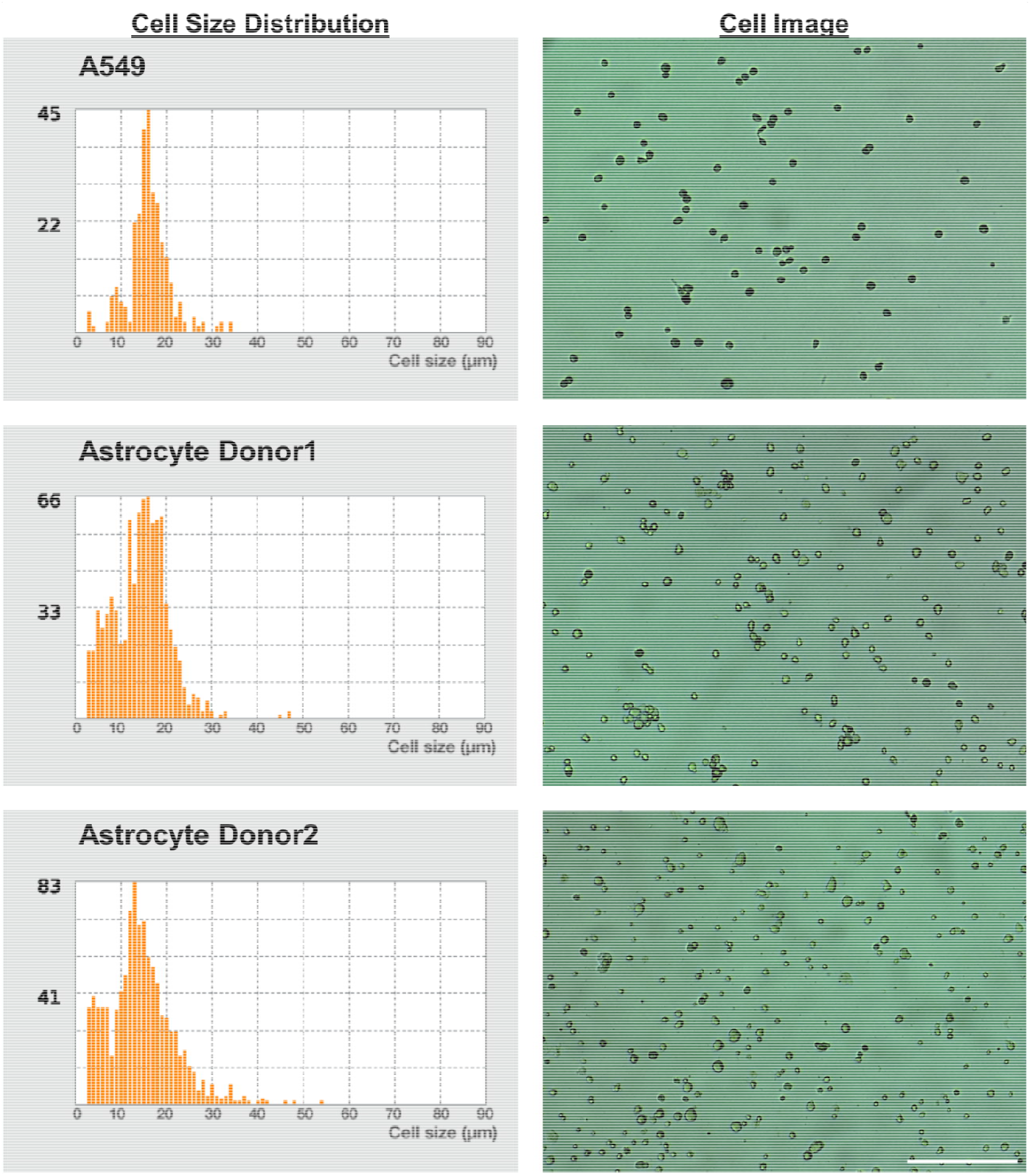
Single cell information prior to dispensing. Example cell size distributions and image of the respective cell suspension of the three cell types tested as captured by the Logos Biosystems Luna-II cell counter. Scale bar represents 500 μm. Average cell size: A549 = 16.9 µm Astrocyte Donor1 = 14.8 µm Astrocyte Donor2 = 15.2 µm

**Fig. S2.**
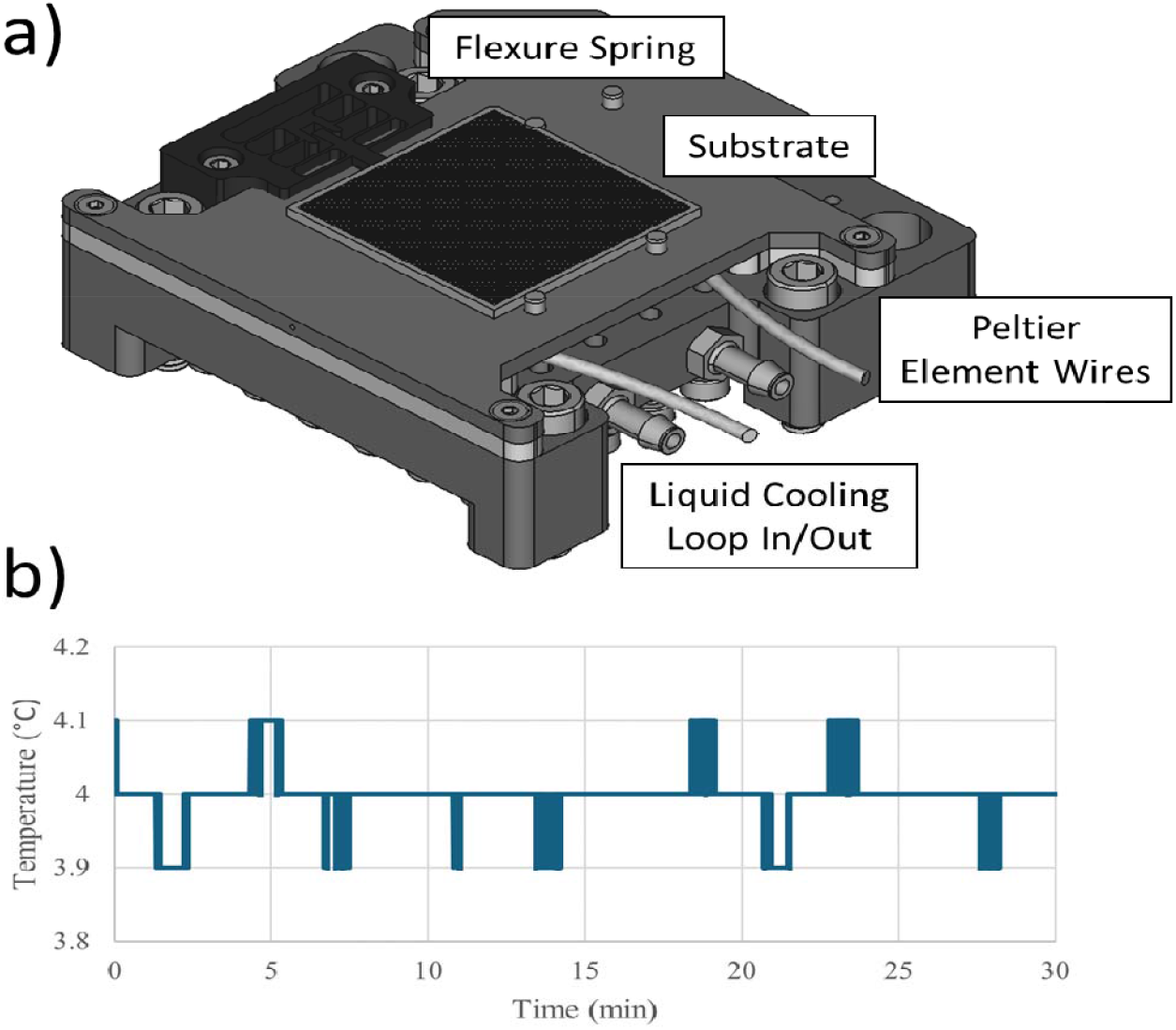
Temperature regulation of Isolatrix. a. The substrate temperature regulation unit. A machined aluminum construction around a Peltier cooling unit. A Delrin middle layer thermally insulates between the hot and cold side of the Peltier element. The substrate is positioned in place via a FDM 3D printed flexure spring. The element is regulated via a liquid cooling loop which continuously pumps room temperature unit across the hot side. An embedded thermistor monitors the temperature of the substrate and provides feedback to the control unit. b. Thermistor reading of the substrate temperature at a setpoint of 4°C over 30 minutes.

**Fig. S3.**
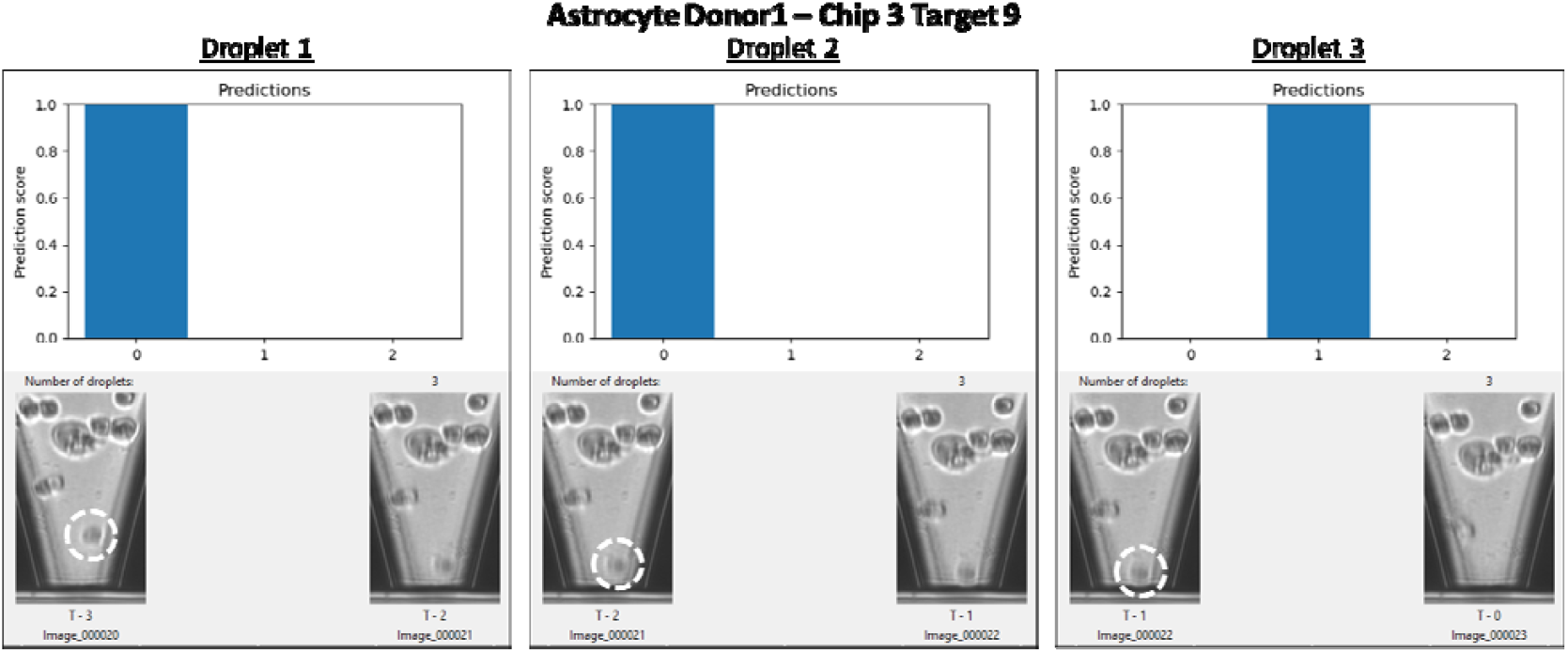
Cell-calling user interface on the Isolatrix. The interface displays a time-series of image pairs and the corresponding neural network output for each droplet deposited into a given well. The neural network assigns a classification score to each class (0 (no cell), 1 (single cell), and 2 (multiple cells)) and the class with the highest score is taken as the predicted outcome. In this example, two consecutive empty droplets were dispensed into the well before a single-cell event was detected. The dispensed cell is highlighted in the pre-dispense image for clarity.

**Fig. S4.**
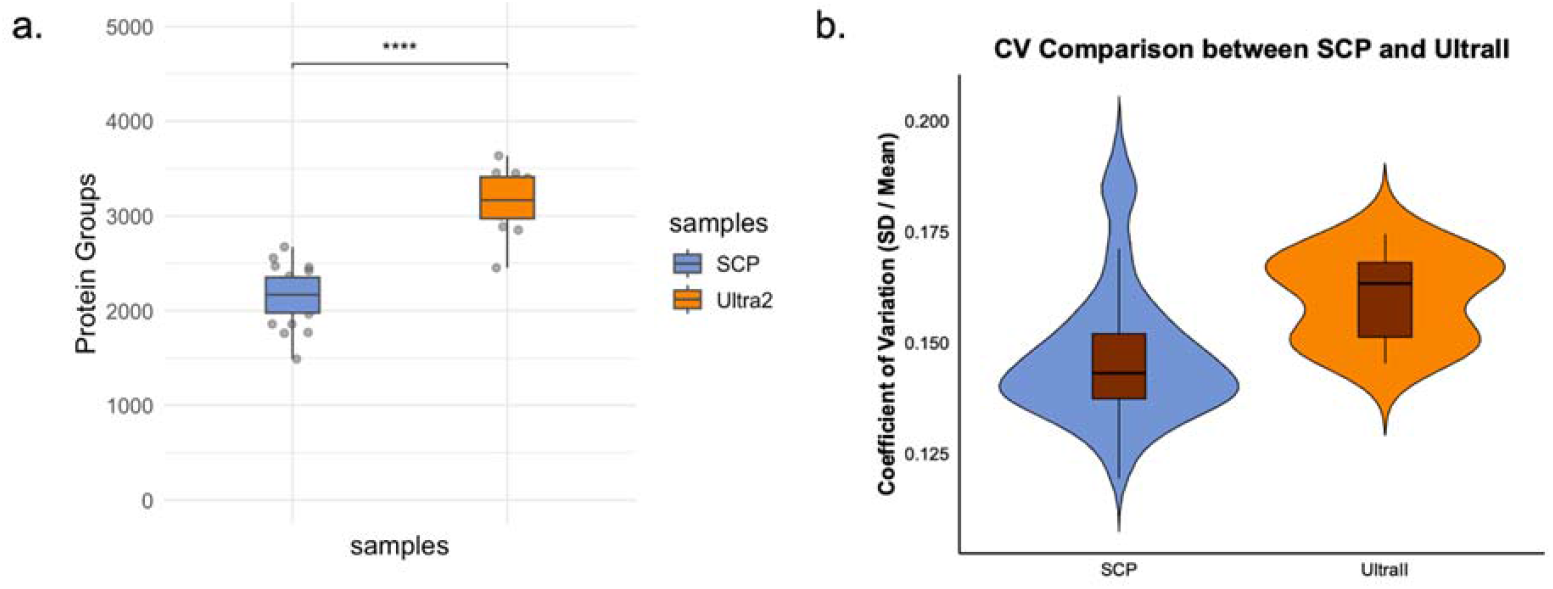
Comparison between timsTOF SCP and Ultra2 data. a. Protein groups identified from A549 injected on timsTOF SCP (blue) and Ultra2 (orange). b. CV comparison between data generated from timsTOF SCP (blue) and Ultra2 (orange).

**Fig. S5.**
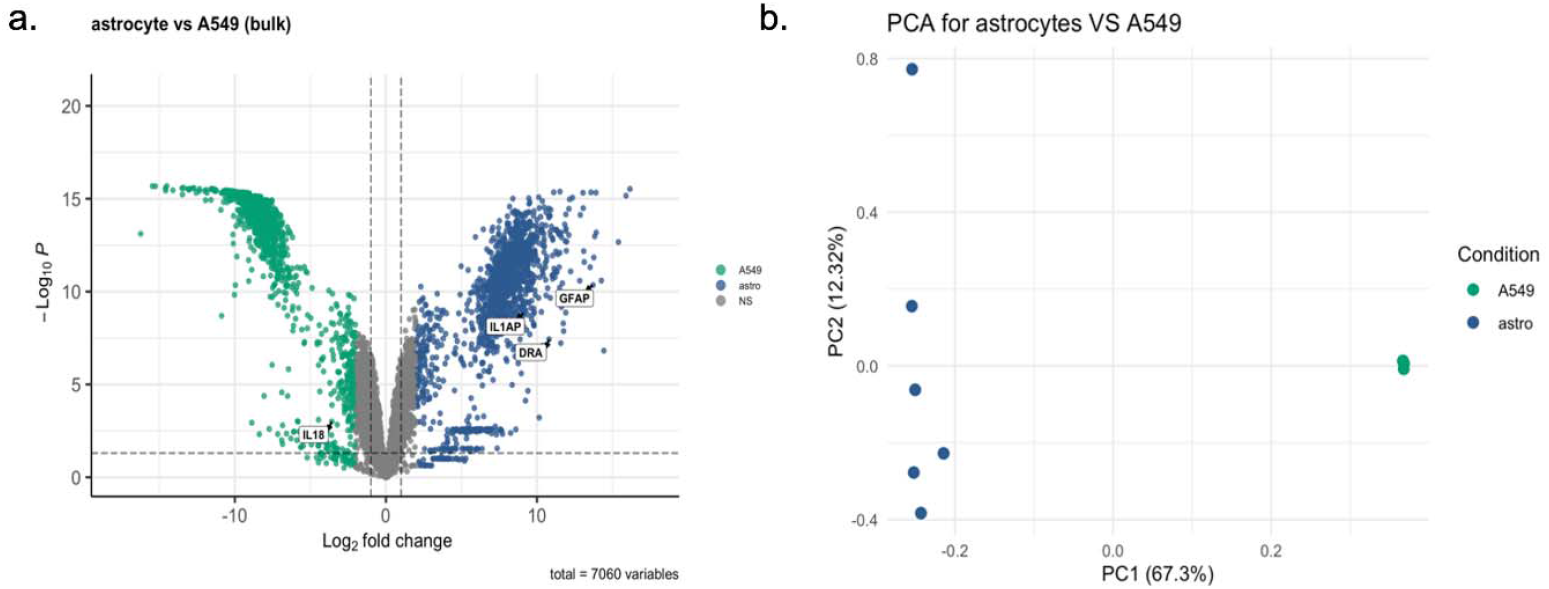
Bulk analysis on A549 compared to primary astrocyte for protein marker identification. Volcano plots on the left showed proteins significantly expressed for A549 in green and primary astrocyte in blue. PCA plot on the right presented clustering for primary astrocyte (including both donors) in blue and A549 in green.

**Fig. S6.**
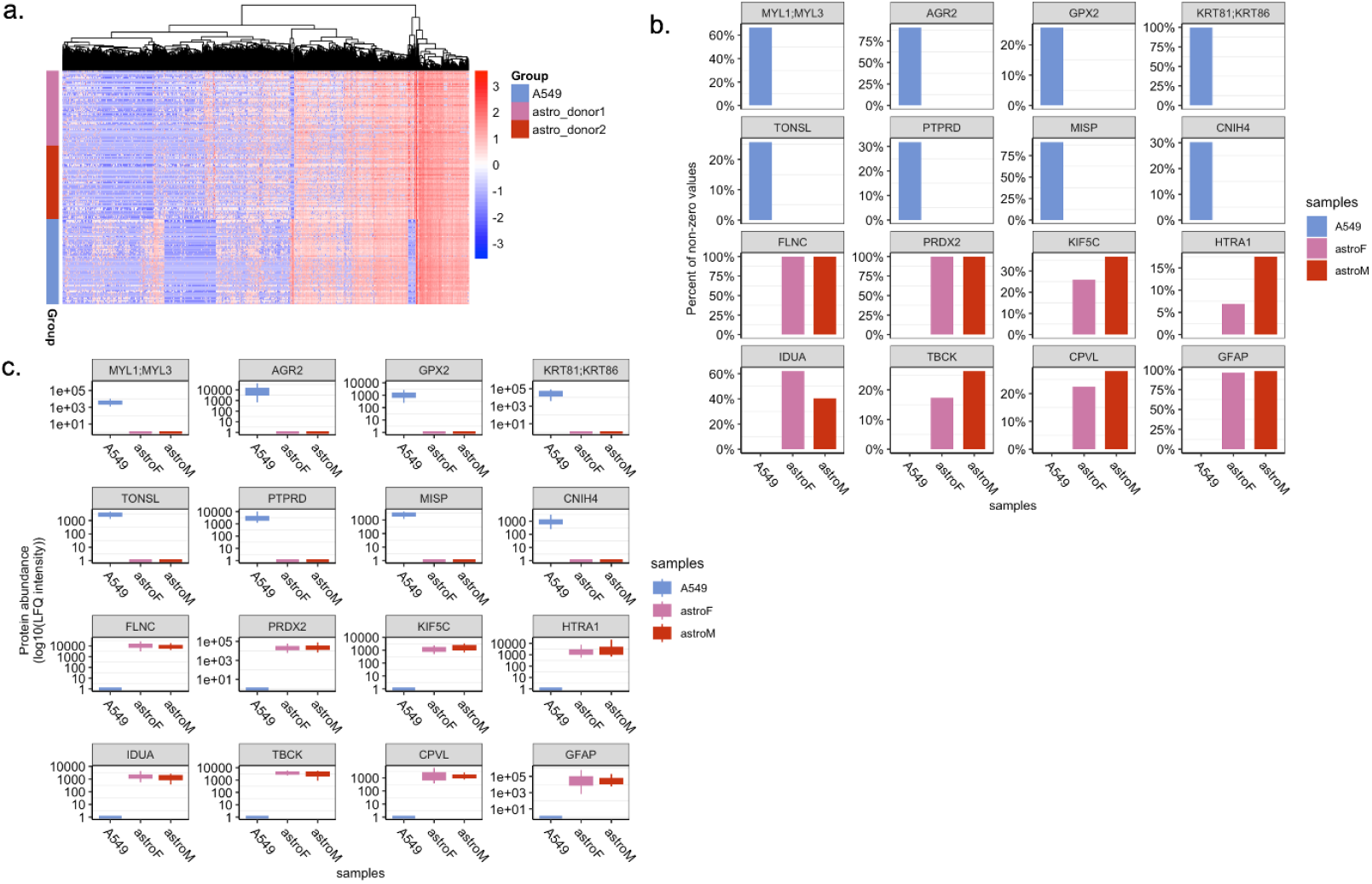
Protein marker comparison between A549 cells and astrocytes. a. Overall heatmap presenting the difference between A549 and astrocyte protein composition. b-c. Proteins identified through limma differential expression analysis were used to evaluate single-cell data quality. The top eight proteins (MYL1, MYL3, AGR3, GPX2, KRT81, KRT86, TONSL, PTPRD, MISP, and CHIN4) represent A549 markers, whereas the bottom eight (FLNC, PRDX2, KIF5C, HTRA1, IDUA, TBCK, CPVL, and GFAP) are astrocyte-specific markers. b. Bar plots showing the protein completeness in A549 and astrocytes. c. Box plots showing the intensity level of the same proteins.

