## Supplementary Information for "SPIN: Inkjet-Driven Nanowell Workflow for Scalable and Sensitive Single-Cell Proteomics"

Eric Cheng *et al.*

**This PDF file includes:**

Supplementary Text

Figs. S1 to S6

Supplementary Text

The supplementary section includes extended technical descriptions of our SPIN set-up and additional data analysis used to support the discussion.

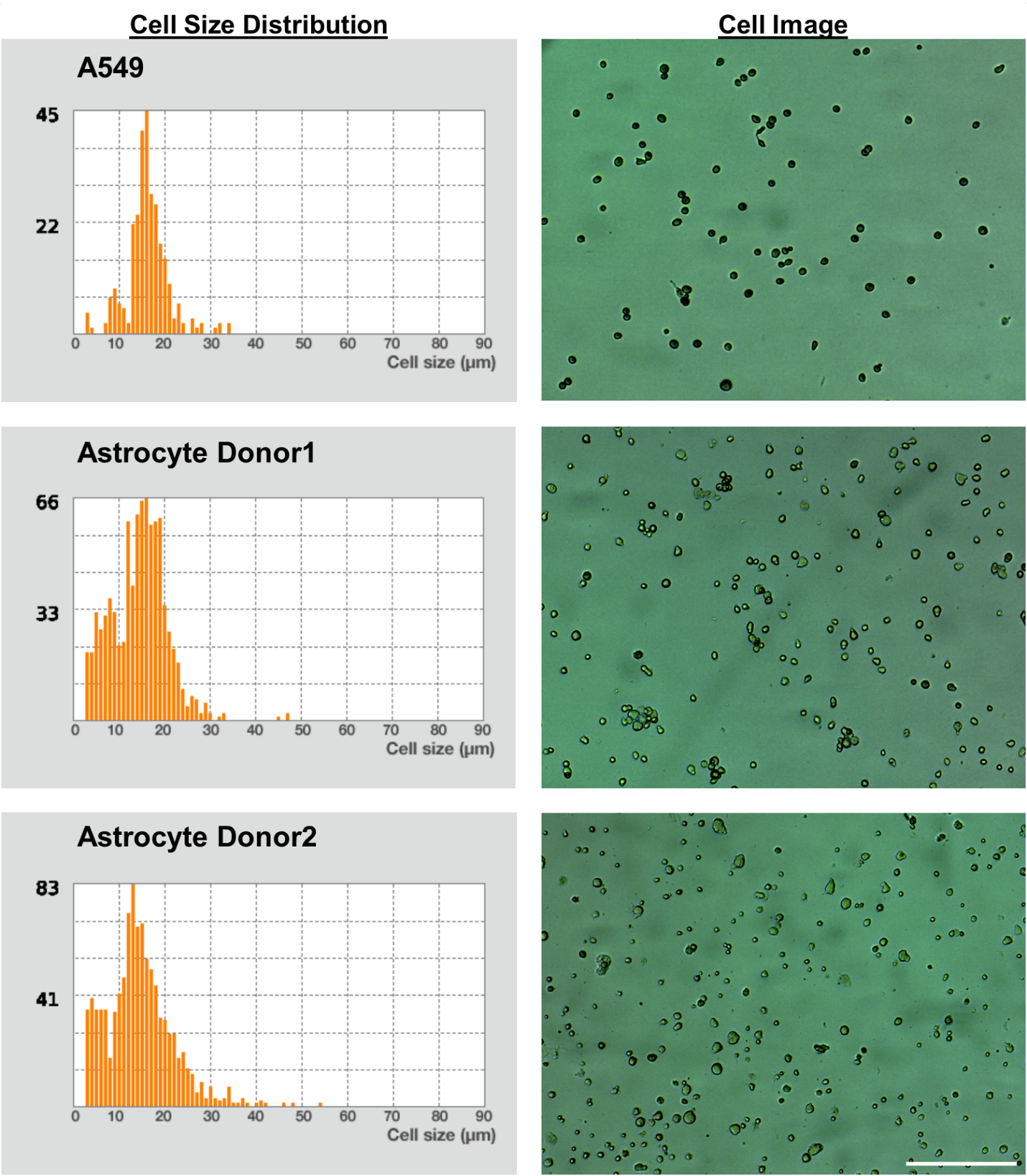
Fig. S1.

Single cell information prior to dispensing. Example cell size distributions and image of the respective cell suspension of the three cell types tested as captured by the Logos Biosystems Luna-II cell counter. Scale bar represents 500 μm.

Average cell size:

A549 = 16.9 µm

Astrocyte Donor1 = 14.8 µm

Astrocyte Donor2 = 15.2 µm

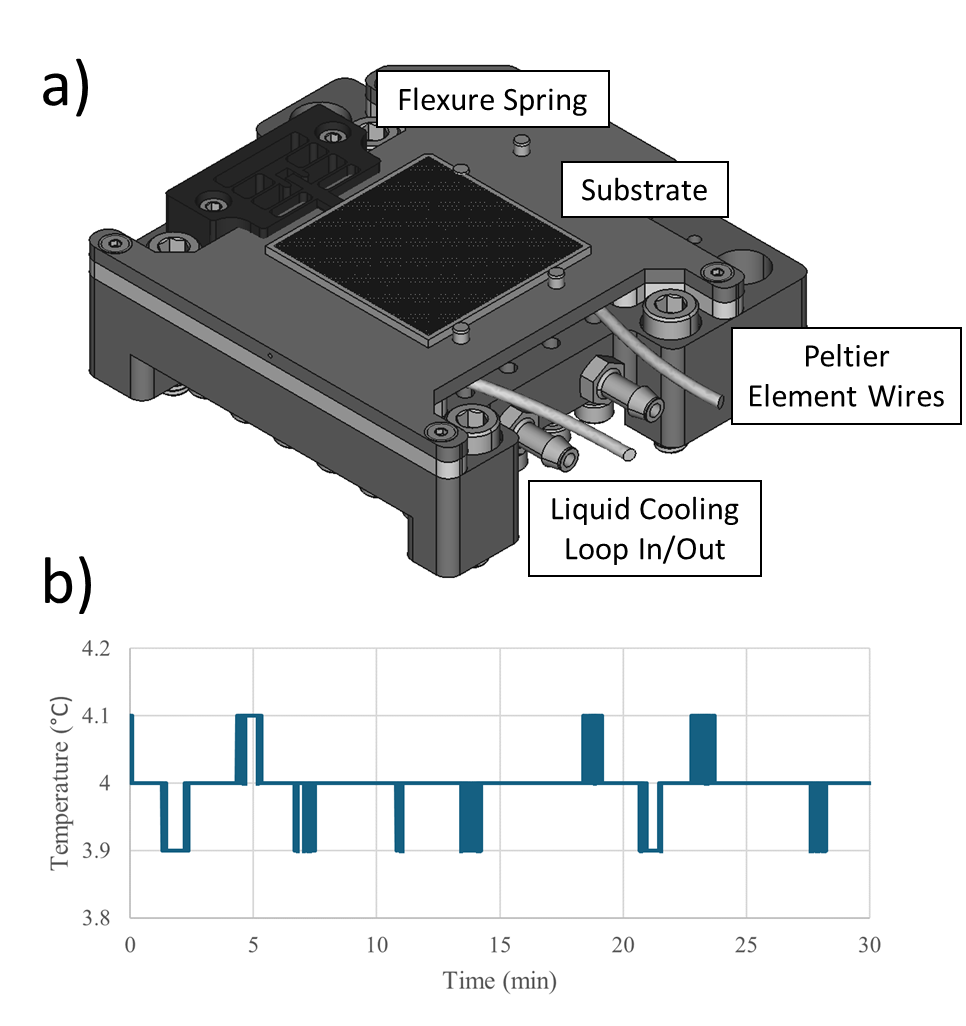

Fig. S2.

Temperature regulation of Isolatrix. a. The substrate temperature regulation unit. A machined aluminum construction around a Peltier cooling unit. A Delrin middle layer thermally insulates between the hot and cold side of the Peltier element. The substrate is positioned in place via a FDM 3D printed flexure spring. The element is regulated via a liquid cooling loop which continuously pumps room temperature unit across the hot side. An embedded thermistor monitors the temperature of the substrate and provides feedback to the control unit. b. Thermistor reading of the substrate temperature at a setpoint of 4°C over 30 minutes.

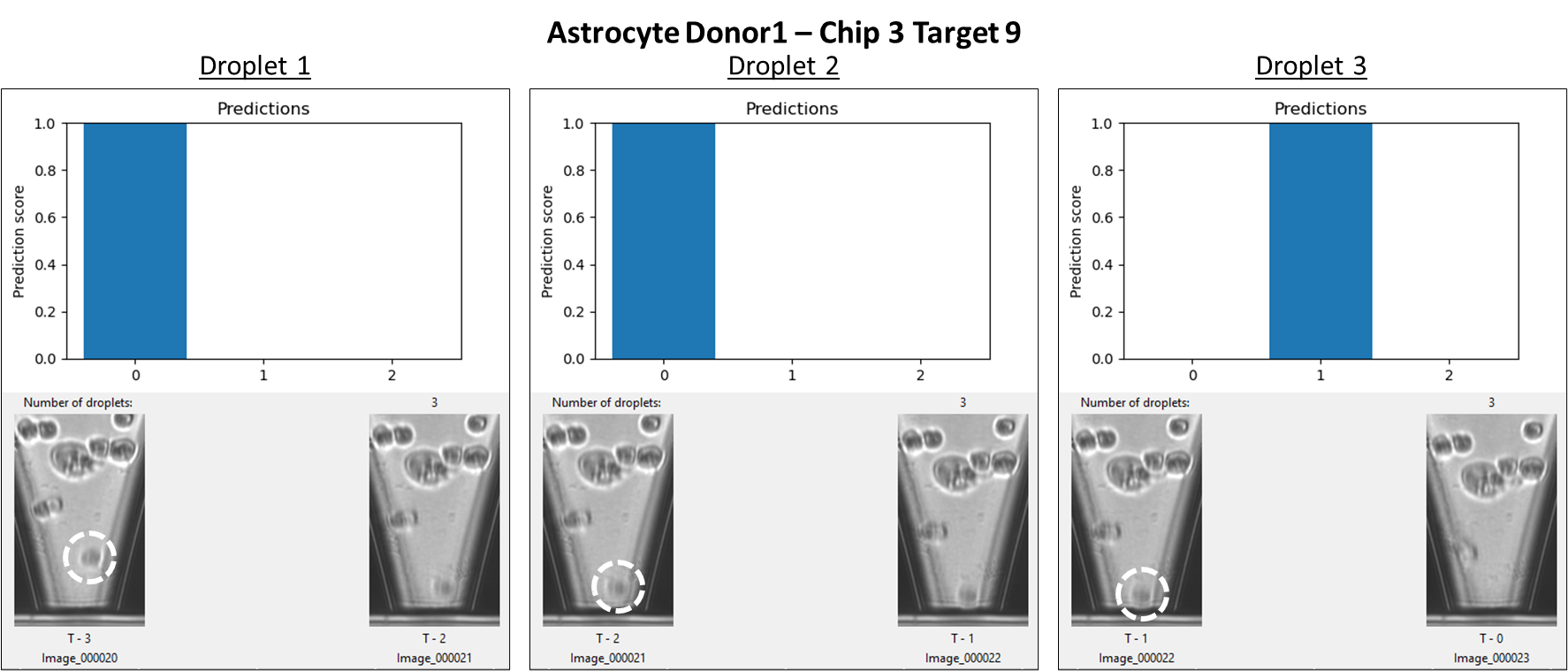

Fig. S3.

Cell-calling user interface on the Isolatrix. The interface displays a time-series of image pairs and the corresponding neural network output for each droplet deposited into a given well. The neural network assigns a classification score to each class (0 (no cell), 1 (single cell), and 2 (multiple cells)) and the class with the highest score is taken as the predicted outcome. In this example, two consecutive empty droplets were dispensed into the well before a single-cell event was detected. The dispensed cell is highlighted in the pre-dispense image for clarity.

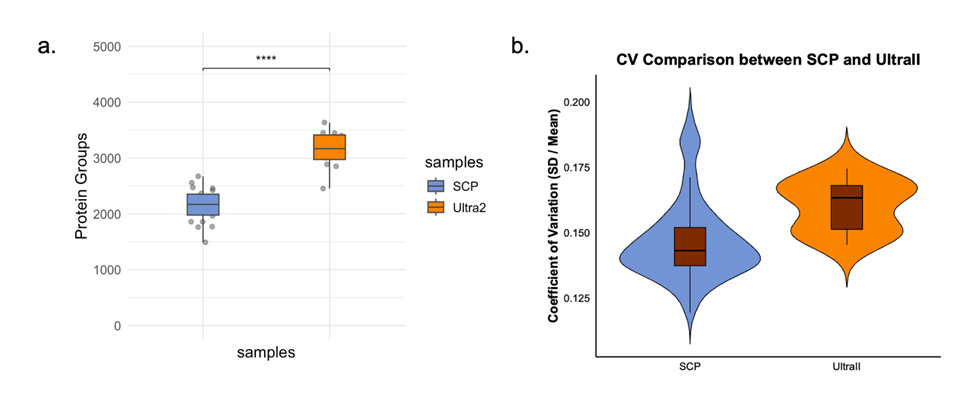

Fig. S4.

Comparison between timsTOF SCP and Ultra2 data. a. Protein groups identified from A549 injected on timsTOF SCP (blue) and Ultra2 (orange). b. CV comparison between data generated from timsTOF SCP (blue) and Ultra2 (orange).

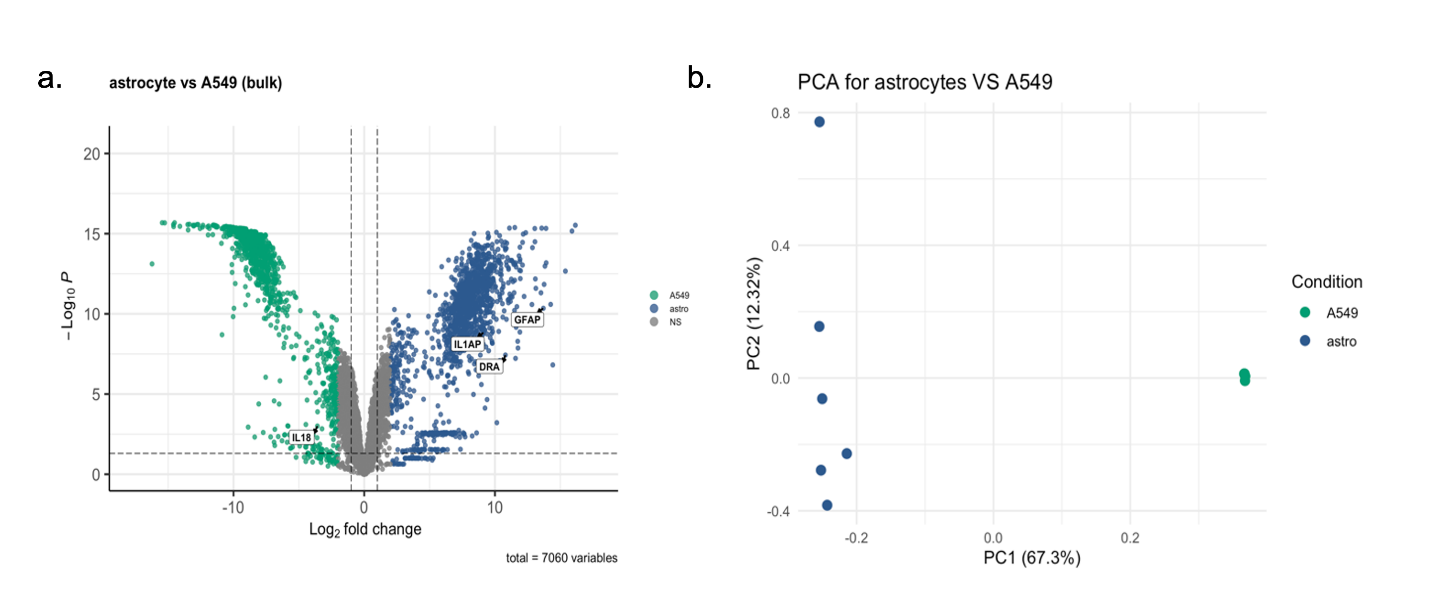

Fig. S5.

Bulk analysis on A549 compared to primary astrocyte for protein marker identification. Volcano plots on the left showed proteins significantly expressed for A549 in green and primary astrocyte in blue. PCA plot on the right presented clustering for primary astrocyte (including both donors) in blue and A549 in green.

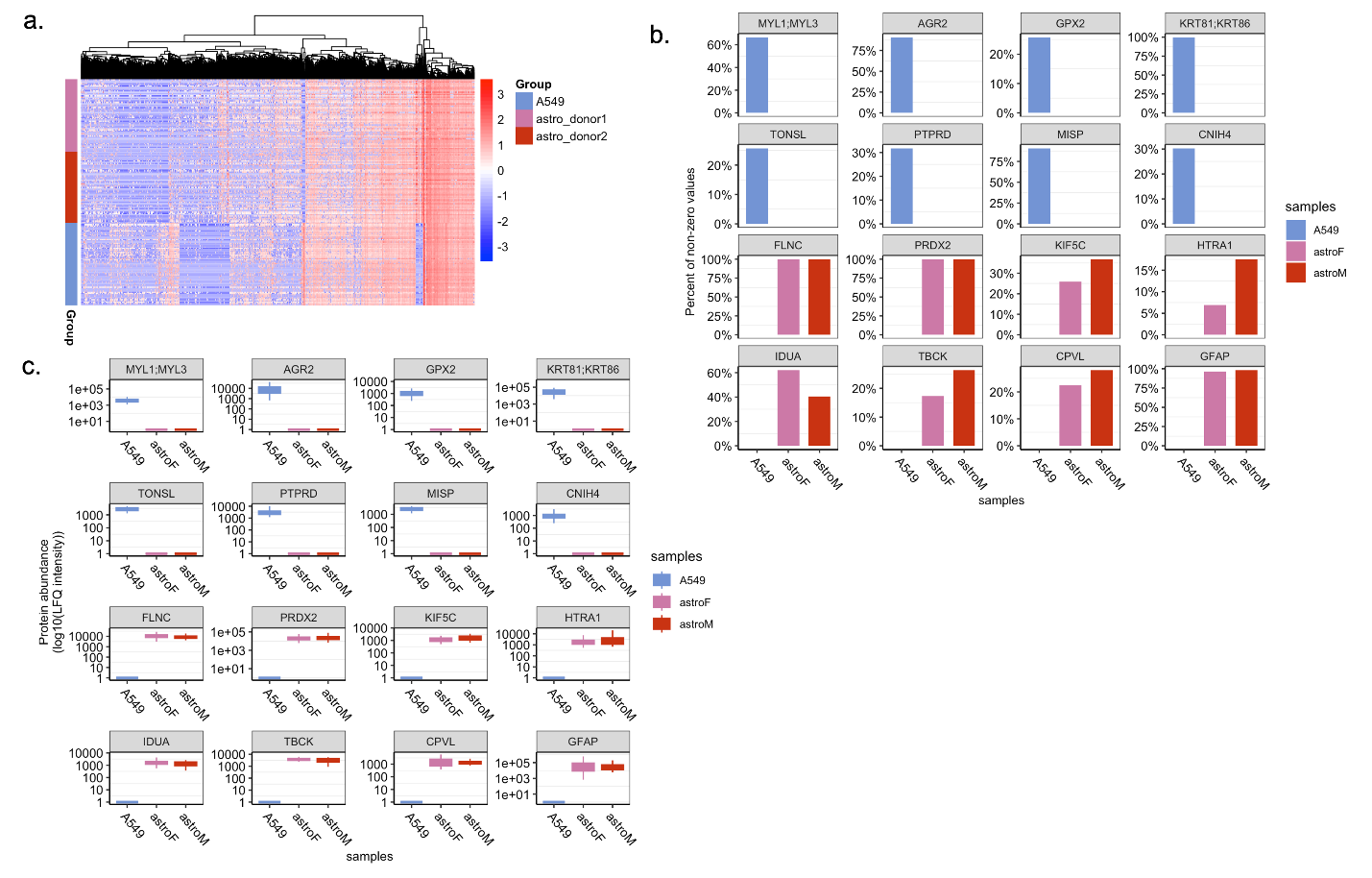

Fig. S6.

Protein marker comparison between A549 cells and astrocytes. a. Overall heatmap presenting the difference between A549 and astrocyte protein composition. b-c. Proteins identified through limma differential expression analysis were used to evaluate single-cell data quality. The top eight proteins (MYL1, MYL3, AGR3, GPX2, KRT81, KRT86, TONSL, PTPRD, MISP, and CHIN4) represent A549 markers, whereas the bottom eight (FLNC, PRDX2, KIF5C, HTRA1, IDUA, TBCK, CPVL, and GFAP) are astrocyte-specific markers. b. Bar plots showing the protein completeness in A549 and astrocytes. c. Box plots showing the intensity level of the same proteins.
